# Open-Source Thermometer, Temperature Controller, and Light Meter for Use in Animal Facilities and During Experiments

**DOI:** 10.1101/2021.05.18.444705

**Authors:** Andrey Andreev, Pavee Vasnarungruengkul, Daniel A. Wagenaar, David A. Prober

## Abstract

Experiments with biological samples require precise control of environmental conditions. In our work we use zebrafish (*Danio rerio*) to understand the neurobiology of sleep, which requires precise control of temperature and lighting. Like many labs, lighting and temperature in the animal facility are centrally controlled in the building. During behavioral experiments and microscopy sessions, we use custom-built heating systems and perform occasional manual checks of conditions. However, without a system to precisely record conditions, gradual changes in temperature can go unnoticed for a long time, and temporary failures may be missed entirely. Here we present the design and characterization of affordable open-source tools to record temperature and light conditions during animal experiments using an Arduino microcontroller or a Raspberry Pi compact computer. The waterproof temperature sensor has high stability over 50 days of recording and is precise to 0.1°C. The Arduino device can be used through a common serial port interface for which we present code in Python and MATLAB. The Raspberry Pi version can be accessed through a web interface, for which we provide an installation guide. We use the device to record and review temperature and lighting conditions in two zebrafish animal facilities. We use our platform to add a water heating system to maintain temperature at 28°C during *in vivo* light-sheet imaging of larval zebrafish. We show that a change in temperature from 28°C to 32°C affects resting heart rate of the animal, highlighting the importance of maintaining and recording conditions. The protocols presented here do not require advanced engineering, fabrication, or software skills, and provide an approach to accurately record and report experimental conditions.

## Introduction

Animal facilities and experiments that use animals require precise monitoring and control of environmental conditions such as light and temperature. This is especially true for sleep studies where experiments can last for several days and environmental conditions affect behavior. Zebrafish and other animals that do not maintain body temperature require control of these conditions by external means. In zebrafish, light and temperature affect several bodily functions such as brain activity, heart rate, and behavior [1–5]. Animal facilities often rely on daily manual data recordings, but this usually only provides a single data point each day and is susceptible to human error. Light measurements are especially important in experiments that involve visual function, where even a small amount of light can perturb rhodopsin, and thus light levels must be continuously monitored during experiments that can last for several weeks [6]. Measurements with automatic continuous sensors provide independent information about the state of the conditions in the animal facility or during an experiment.

Tracking conditions and independent verification of set points and schedules such as cyclical light conditions allows more precise experiments as well as reporting of the actual data, rather than estimates and assumptions about the conditions. Commercially available thermometers with data logging can cost up to $250, require specialized software, and usually cannot have additional sensors or interface for remote access. Here we present an open-source platform for a temperature and light data logger with several options for the interface, including remote web access. Device cost starts at $40. This platform can also be used to regulate temperature in a sample chamber using a simple heating element and proportional control scheme, similar to previously reported devices to keep and rapidly change temperature conditions under a stereomicroscope [7] or confocal microscope [8].

## Results

Our temperature recorder (Figure 1A) consists of a microcontroller (Arduino Uno, see Table 1 for list of parts) and a waterproof digital temperature sensor (DS18B20). Assembly consists of the five parts listed in Table 1 soldered onto a “shield” (general purpose soldering circuit board). The Arduino code for measuring temperature and light levels is presented in Code Snippet 1.

**Figure 1.**
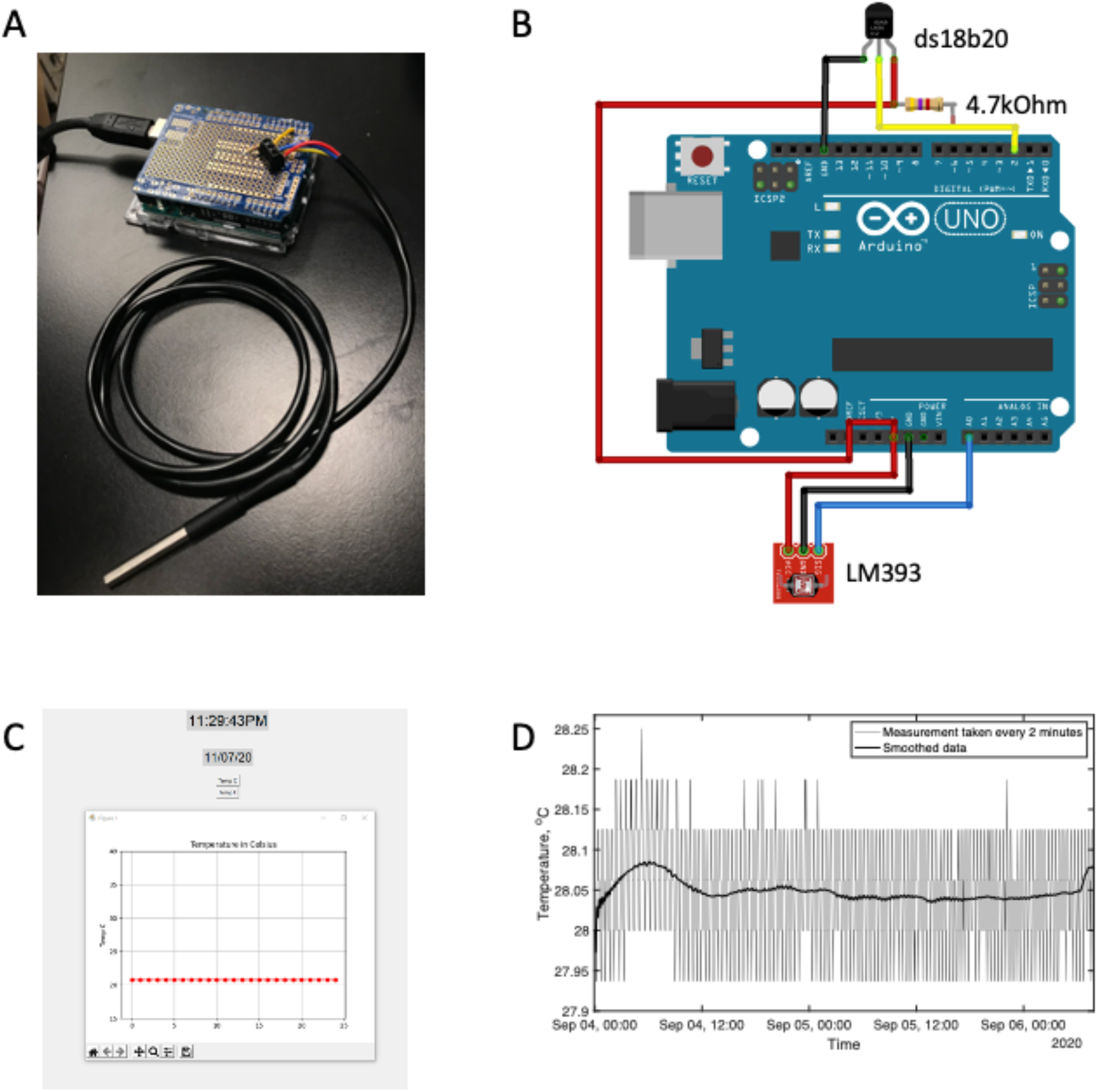
Arduino device for measuring temperature and associated interface allows long-term recording with a PC interface. (**A**) Arduino Uno microcontroller with waterproof temperature sensor DS18B20 in a steel jacket. (**B**) Wire diagram of Arduino Uno with temperature sensor and light detector attached. The light detector provides analog output that is connected to an analog input of the microcontroller (blue wire), 5V (red wire), and ground (black wire). The temperature sensor DS18B20 uses a 1-wire protocol and is connected to ground (black wire), and a digital pin 2 (yellow wire) with a pull-up resistor to 5V (red wire). (**C**) Screenshot of Python graphical user interface used to read out temperature measurements using a computer. (**D**) Continuous recording of water temperature with 2-minute resolution for over 48 hours. Set point was 28°C. The thick line shows smoothed data (sliding window over 200 measurements).

**Table 1.**
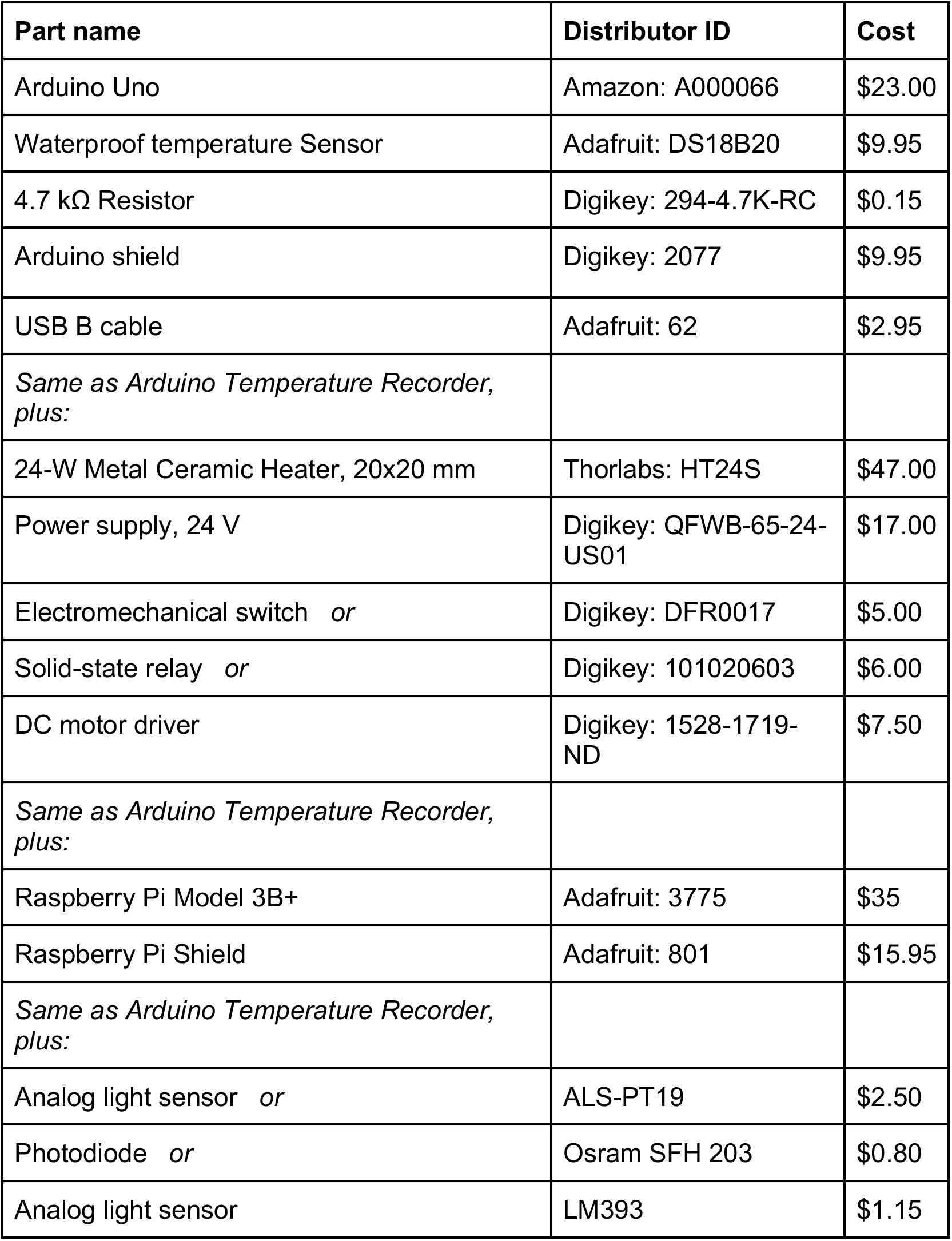
List of parts.

The Arduino Uno device records temperature measurements from the sensor and returns these via a USB port (Figure 1B) at a preset rate between once per second to once every few minutes. To collect and analyze the data, we developed a custom graphical user interface using Python (Figure 1C, Code Snippet 2-1 and 2-2), and MATLAB (Figure 1D, Code Snippet 3). We first tested the sensor by recording temperature in a high-precision water bath set at 28°C. Continuous recording for 48 hours (Figure 1E) yielded temperature measurements of 28.04°C±0.06°C (mean ± SD). We did not determine whether this variation comes from the heater or the sensor. Thus the sensor is stable within <0.1°C, and is more precise than a previously reported similar thermometer [8].

The open Arduino platform allows use of multiple types of sensors. The photodiode (Osram SFH 203, Table 1) that produces current as a function of illumination intensity can be directly connected to analog input to provide measurement of the ambient light level. Continuously monitoring light level or temperature can identify irregularities due to unscheduled entrances by facilities personnel. We have also tested an analog sensor (ALS-PT19, Table 1) that is calibrated to reflect the human visual spectrum, and a photoresistor (LM393, Table 1).

Arduino microcontrollers do not directly support internet connectivity that is necessary for remote access to the data. However, they can be combined with an inexpensive and compact Raspberry Pi computer. Using common functions, we implemented an open-source web interface (Figure 2A) that allows access to raw recorded files, as well as interactive plotting of the data (Figure 2B). One scenario where this device was useful for us is a newly established animal facility. Tracking for more than two weeks identified unexpected variations in temperature control and lighting conditions, and provided concrete data that could be used to determine the source of the variations. We collected data on two different animal facilities, before and after the move to a new building (Supplemental Figure 1).

**Figure 2.**
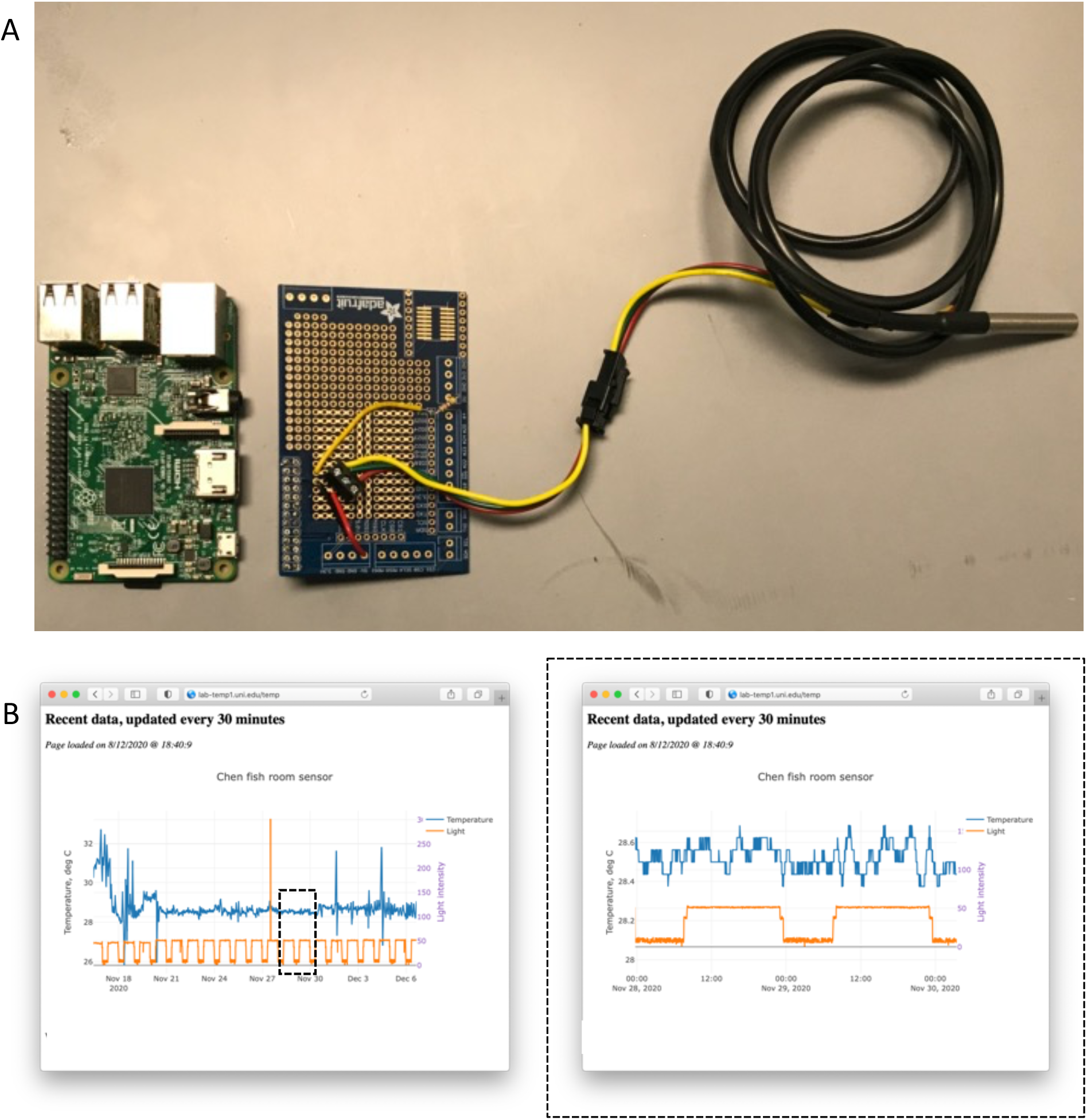
Connection of a temperature sensor to a Raspberry Pi computer allows remote monitoring of temperature and light levels with 2-minute resolution in an animal facility. (**A**) Photograph of Raspberry Pi 3B+ and a corresponding shield with installed DS18B20 water-proof temperature sensor. Green wire connects to ground. Red wire connects to 5V. Yellow wire connects to IO4 pin through 4.7kΩ pull-down resistor. (**B**) Screenshot of the web interface of conditions recorded over 20 days in a zebrafish animal facility. Temperature and light intensity are reported and the user can interactively expand a region of interest (e.g., dashed box in left panel is enlarged in right panel) and save the plots as an image.

With a single Arduino device, a temperature sensor can be bundled together with a heater to create a control device to keep conditions at a given temperature during microscopy sessions (Figure 3A). We applied a simple non-linear proportional control process (Code Snippet 5) to regulate the temperature of a 50-mL imaging sample chamber (40 mm x 60 mm x 20 mm, Figure 3A) in a custom light-sheet microscope similar to a previously published design [9]. We used a commercially available ceramic heating element and the thermometer described in the previous section (Arduino-based thermometer). This device brought the temperature within the sample chamber to 28°C from a room temperature of 21°C in less than 5 minutes (Figure 3D). The sample temperature was maintained at 28°C ± 0.3°C. We verified temperature values using second calibrated thermometer (Figure 3B). We used Arduino code to read the temperature (*T*), and if it fell below the target setpoint (*T*_ref_), the relay was switched on, closing the circuit and turning the heater on for a given period of time. We found that fast and accurate control is achieved if the minimal time of heater activation is set to be 10 ms, with an additional 100 ms per degree of temperature difference between the reading and the setpoint:

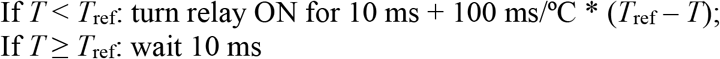

**Figure 3.**
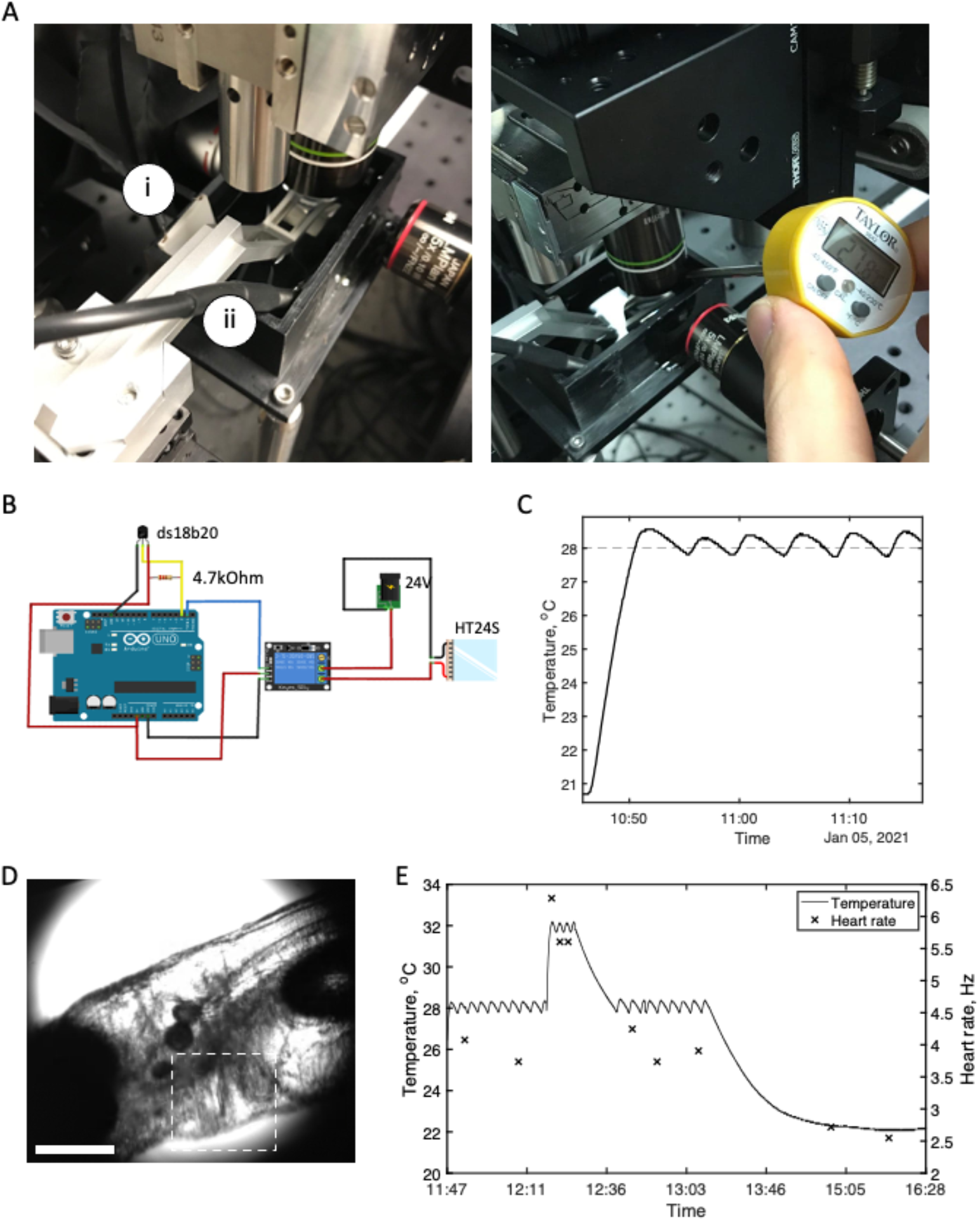
The Arduino temperature sensor can be modified and programmed to drive proportional control of temperature inside a sample chamber. (**A**) Assembled device attached to the sample chamber of a custom light-sheet microscope. (i) Heating element (ceramic plate) and (ii) thermometer are submerged in water. Recorded temperature deviated from independent lab thermometer (shown here) by less than 0.3°C. (**B**) Wiring diagram of the heating device. Thermometer DS18B20 is attached as described in Figure 1B. In addition, the electromechanical relay is connected to 5V (red wire) and ground (black wire) and activated by a digital signal from pin 1 (blue wire). Activation of the relay causes the connection of the heating element to a 24 V power source. By regulating the duration of relay activation, the amount of heat that is added to the chamber is controlled. (**C**) Ramp of temperature in the chamber from a room temperature of 23°C to 28°C takes less than 5 minutes and temperature is subsequently maintained within ±0.5°C. The target temperature is indicated by the dashed line. (**D**) 5 day old zebrafish (5 dpf) whose heart is imaged using transmitted light. The heart region is indicated with a dashed box, which outlines the region analyzed using Fourier transformation to extract frequency of image change. Scale bar: 200 μm. (**E**) Simultaneous recording of temperature in the sample chamber and larval zebrafish heart rate. The temperature was first maintained at 28°C, then stepped up to 32°C, then allowed to decrease to 28°C, and subsequently to room temperature (22°C). Heart rate was recorded as described in Supplemental Figure 2. Without active cooling, it took ~10 minutes for the temperature to decrease from 32°C to 28°C.

Heating a 50 mL water chamber from 28°C to 32°C took less than 1 minute (Figure 3D). Return to 28°C took 10 minutes without any active cooling device with a room temperature of 22°C. We observed effects of temperature on zebrafish heart rate using imaging at different temperatures. We used a behavioral camera on our light-sheet microscopy setup to register heart movement, imaging at 40 frames per second (Supplemental Figure 2). Extraction of heart rate from the imaging data was done using an image difference approach and a Fast Fourier Transform. Heart rate correlated with temperature as previously described [10], changing from 4 Hz at 28°C to 6 Hz at 32°C, and then to 2.5 Hz at 25°C (Figure 3F).

## Discussion

We described an open-source temperature recording and control scheme based on Arduino microcontroller architecture using commodity parts and open-source software. We showed how this platform can be expanded to provide long-term recording of light and temperature conditions in an animal facility. Furthermore, an Arduino platform can be used as a basis for a custom temperature control device to keep a light-sheet microscope sample chamber at a constant temperature.

Accurate reporting of temperature is important during animal experiments. By specification, the DS18B20 thermometer has ±0.5°C accuracy, and might need calibration. We have not done extensive calibration, but temperatures reported by the DS18B20 sensor consistently agreed with other thermometers to within 0.5°C.

Open-source hardware allows expansion of the set of sensors. This allows continuous collection of values of multiple environmental parameters, unlike the usual practice of occasional checks. Temperature and humidity can be monitored using a single sensor (e.g., DHT22). A vibration sensor (using an analog accelerometer such as ADXL335) can be used to record and validate a vibration-based assay, such as tapping a plate containing zebrafish larvae, which can be used as an arousal threshold assay [11]. A microphone can be installed for experiments that involve sound stimulation. Sensors can be combined according to the needs of a particular experiment to provide continuous recording and calibration of the assay, as well as provide data for reporting experimental methods.

The sample chamber temperature described here used a simple proportional control algorithm. It can be improved by adding a power-regulator (e.g., using a motor driver) and adding integrated and differential control. Heating elements and thermometers come in different form-factors (such as rods, circular elements, or flexible films), and potentially can be adapted for various microscopes, imaging chambers, or experimental platforms, without changing the principle of operation.

Attaching sensors to a computer allows more external software to be run, such as email alerts and remote access to data on a network. We integrated a simple web interface to observe temperature and light conditions in an animal facility, but issues of security and continuous operation of the computer are often not trivial. We coordinated with our campus to block all incoming internet access to our device, only allowing access via a virtual private network. We also disabled access using passwords, relying only on public/private key pair authentication. Attaching simple devices to outdated campus networks can potentially compromise IT security.

The design and approach presented here allows continuous automatic reporting of environmental data, and calibration of a wide range of experimental setups that can result in decreased experimental variability. We hope that open-source devices such as the device presented here will enable scientists to track and report data about experimental conditions in their publications.

## Methods

The DS18B20 waterproof temperature sensor uses a OneWire protocol [12] that requires only one data pin in addition to a power (5V) and ground connection. In order to create the Arduino-based temperature and light recorder, we connected the DS18B20 sensor probe onto the Arduino shield. As shown in Figure 1A, the yellow data wire was linked to digital port 2, the red wire to 5V, and the blue wire to GND. The OneWire protocol requires the use of a 4.7-kΩ pullup resistor that connects the data wire of the sensor to the 5V wire. For the Raspberry Pi version of the device, the data wire was connected to IO4 pin, otherwise wiring is unchanged. We followed a tutorial accessible at https://bit.ly/3u5e0uJ.

To record ambient light levels, we used a LM393 photo-resistive light sensor. Similar to the temperature probe, the LM393 photo-resistive light sensor has 3 wires, COM, 5V, and GND. The COM wire was connected to the Arduino A0 analog input pin, the 5V wire was connected to the board 5V pin, and the GND wire was connected to the ground pin.

Another light sensor that we tested, ALS-PT19, is an analog RoHS-compliant sensor with ground, 3–5 V, and data pins. It was wired in the same way as LM393.

Wire diagrams were created using Fritzing [13]. We used MATLAB versions 2018b and 2020b, and Python version 3.1. We tested Python and MATLAB code on Windows 7, Windows 10, and Mac OS X 10.15.

The Raspberry Pi version of the thermometer/light-meter was assembled similarly to the Arduino version. The source code is available at https://github.com/aandreev0/raspb-temp-logger. The installation process consists of these steps:

1. Install recent version of Raspbian operating system
2. Connect to the Raspberry Pi using an external display and keyboard
3. Get the MAC address and communicate with the IT services of your institution to allow access via a separate domain name, for example “`lab-thermo1.uni.edu`”.

a. MAC address is shown on the start-up screen or using the ‘`ifconfig`’ command and looking for line ‘`HWaddr: AA:BB:CC:DD:EE:FF`’ ~~~
[pi@retropie:~ $ ifconfig
eth0       Link encap:Ethernet  HWaddr b8:27:eb:b6:65:eb
           inet addr:131.215.52.121  Bcast:131.215.52.255  Mask:255.255.255.0
           inet6 addr: fe80::5fbl:e273:761:9afe/64 Scope:Link
           UP BROADCAST  RUNNING MULTICAST  MTU:1500  Metric:l
           RX packets:7434728 errors:0 dropped:7724 overruns:0 frame:0
           TX packets:I76005 errors:0 dropped:0 overruns:0 carrier:0
           collisions:0 txqueuelen:l000
           RX bytes:5966I46I3 (568.9 MiB)  TX bytes:I9I593702 (182.7 MiB)
~~~
4. On a personal or lab workstation configure password-less SSH authentication using a private/public key pair:

a. Follow the guide for generation of a private/public key pair on your system:

i. On Linux-type systems and Mac OS X use “`ssh-keygen -t rsa`” command
ii. On Windows systems use PuTTy or OpenSSH, and follow instructions for the corresponding terminal
b. Add a public key to Raspberry Pi’s `~/.ssh/authorized_keys` file
5. Connect the Raspberry Pi to the internet using an ethernet cable.
6. Continue working directly in the Raspberry Pi terminal or connect to it using SSH (step 4).
7. Copy code from GitHub repository to `~/raspb-temp-logger` using “git clone https://github.com/aandreev0/raspb-temp-logger.git”.
8. Install the necessary Python packages using the “`pip install`” command.
9. Install Nginx server using “`sudo apt install nginx`”.
10. Configure Nginx server to serve the content of `~/raspb-temp-logger`:

a. Edit the Nginx configuration file using “`sudo nano /etc/nginx/sites-available/default`”
b. Change the line “`root /var/www/html/;`” to “`root /home/raspb-temp-logger/`”
c. Save and close the file using CTRL+X
d. Restart Nginx using the command “`sudo systemctl restart nginx`”
11. Configure crontab to automatically run the temperature-recording script every 2 minutes using “`sudo crontab -e`”

a. Add line: “`*/2 * * * * python /home/pi/raspb-temp-logger/report_temp.py`”
12. Configure crontab to automatically run the generation of plots every 30 minutes using the same command

a. Add line “`*/30 * * * * python /home/pi/raspb-temp-logger/csv_to_json.py`”
b. Close crontab by pressing CTRL+X
13. Test by accessing the domain name or IP address

To generate plots, we used the open-source Python library *plotly* and custom code that generates a javascript data structure file (JSON file) from CSV files with recorded data.

The light meters we used are analog, with output voltage between 0V and 5V. The Raspberry Pi 3B+ lacks a built-in analog-to-digital converter (ADC) and cannot work with these devices directly. We therefore used an Arduino as an ADC. The light-measuring device was plugged into the analog input of Arduino Uno, and the Arduino was connected to a Raspberry Pi through a USB cable. We used similar code to write light measurement to serial port, and read serial port using Python script (Code Snippet 6).

Heart rate recording was performed on larval zebrafish at 5 days post fertilization by imaging using infrared trans-illumination with an LED (780nm, Thorlabs) and a CMOS camera at 40 frames per second. We used an air 4x objective with 0.1 NA (Edmund Optics). To process the video, we first calculated image difference between adjacent frames (temporal differentiation), followed by a Fast Fourier Transform of average intensity within a region of interest containing the heart (see Supplemental Figure 2 for details). Code of the plugin for Fiji image analysis software is available at https://github.com/aandreev0/fiji-heartrate.

## Acknowledgements

We thank Prober lab members for providing animals for the experiments, and especially Dr. Amina Kinkhabwala for testing the design and instructions. The first version of the code used to extract heart rate from images was developed by A.A. during work in Truong/Fraser lab at University of Southern California. This work was supported by grants from the NIH to D.A.P. (R35 NS122172 and R01 MH121601).

## Code availability

- Code for the Raspberry Pi-based web interface is available at https://github.com/aandreev0/raspb-temp-logger
- Code of the heart rate extraction plugin for Fiji is available at https://github.com/aandreev0/fiji-heartrate

## Code

### Code snippet 1: Arduino interaction with DS18B20 and analog light sensor (LM393)

~~~
void setup(void)
{
  Serial.begin(9600);
  sensors.begin();
}
void loop(void)
{
  sensors.requestTemperatures();
  byte lightValue = analogRead(0); // Read light value from port A0
  Serial.print(sensors.getTempCByIndex(0)); // Read temperature value
  Serial.print(“,”);
  Serial.println(lightValue);
  delay(800); // wait 800ms
}
~~~

### Code snippet 2-1: Python code to get recording via serial port on PC

~~~
import serial
import serial.tools.list_ports
def findArduino(ports):
    commPort = ‘None’
    numPorts = len(ports)
    for i in range(0, numPorts):
        strPort = str(ports[i])
        if ‘Arduino’ in strPort:
            splitPort = strPort.split(‘ ’)
            commPort = (splitPort[0])
    return commPort
ports = serial.tools.list_ports.comports()
connectPort = findArduino(ports)
if connectPort != ‘None’:
    s = serial.Serial(connectPort,baudrate= 9600)
    print (‘Connected to ’ + connectPort)
else:
    print (‘Can\’t find Arduino device’)
dataStr = s.readline().decode(“utf-8”)
dataArray = dataStr.split(‘,’)
temperature = float(dataArray[0])
light       = float(dataArray[1])
print([temperature, light])
s.close()
~~~

### Code snippet 2-2: Python code to make GUI with data plot

~~~
def drawn():
    count = 0
    while True:
        arduinoString = s.readline().decode(“utf-8”)
        dataArray = arduinoString.split(‘,’)
        temp = float(dataArray[0])
        L = float (dataArray[1])
        tempC.append(temp)
        light_sensor.append(L)
        drawnow(makeFig)
        plt.pause(.000001)
        count = count + 1
        if (count > 50):
            tempC.pop(0)
            light_sensor.pop(0)
~~~

### Code snippet 3: MATLAB code to get recording via serial port

~~~
port = ‘COM4’;
s = serial(port, ‘Terminator’,‘LF’);
fopen(s)
getString = fgets(s);
data = regexp(getString, ‘,’, ‘split’);
temperature = str2num(data{1}) % temperature in C
light       = str2num(data{2}) % light in a.u.
fclose(s)
~~~

### Code snippet 4: Python code to read temperature from Raspberry Pi

~~~
import os
import glob
import sys
import serial
os.system(‘modprobe w1-gpio’)
os.system(‘modprobe w1-therm’)
light_arduino = ‘/dev/’
def read_temp_raw():
    base_dir = ‘/sys/bus/w1/devices/’
    devs = glob.glob(base_dir + ‘28*’)
    if len(devs) > 0:
        device_folder = devs[0] # only picks 1st device
        device_file = device_folder + ‘/w1_slave’
        f = open(device_file, ‘r’)
        lines = f.readlines()
        f.close()
        return lines
    else:
        return False
def read_temp():
    lines = read_temp_raw()
    if not lines:
        return 0
    while lines[0].strip()[-3:] != ‘YES’:
        time.sleep(0.2)
        lines = read_temp_raw()
    equals_pos = lines[1].find(‘t=’)
    if equals_pos != −1:
        temp_string = lines[1][equals_pos+2:]
        temp_c = float(temp_string) / 1000.0
        temp_f = temp_c * 9.0 / 5.0 + 32.0
        return temp_c
~~~

### Code snippet 5: Arduino code for the proportional control of sample chamber heater

~~~
#include <OneWire.h>
#include <DallasTemperature.h>
#define ONE_WIRE_BUS 2 // DS18B20 COM is plugged into pin A2
int relayPin = 5;      // digital pin used to switch relay on/off
OneWire oneWire(ONE_WIRE_BUS);
DallasTemperature sensors(&oneWire);
float temp;
float set_temp = 28.5; // target temperature
int relayStatus; // status of the switch: 1=open, 0=closed
void setup(void)
{
  Serial.begin(9600);
  sensors.begin();
  pinMode(relayPin,OUTPUT);
  digitalWrite(relayPin, LOW);
  relayStatus = 0;
}
void loop(void)
{
  sensors.requestTemperatures(); // get temperature reading
  temp = float(sensors.getTempCByIndex(0));
  // print current time, current remperature, and relay position
  Serial.println(String(millis())+“:”+String(temp)+“,”+relayStatus);
  if(temp < set_temp){
      digitalWrite(relayPin,HIGH);        // turn heater ON
      relayStatus = 1;
      delay(10 + int((set_temp - temp)*100)); // wait 10ms+100ms/deg C
  }else{
      digitalWrite(relayPin,LOW);        // turn heater OFF
      relayStatus = 0;
      delay(10);
  }
}
~~~

### Code snippet 6: Python code for light measurement from serial port in Raspberry Pi

~~~
import serial
def read_light():
    ser = serial.Serial(‘/dev/ttyACM0’)  # open serial port
    avg_light = 0;
    for i in range(0,5):
        line = ser.readline()  # read a ‘\n’ terminated line
        if i>2:
            avg_light += int(line.strip())
    avg_light = avg_light / 2.0;
    ser.close()
    print(“Averaged light:” + str(avg_light))
    return avg_light
~~~

## Supplemental Figures

**Supplemental Figure 1.**
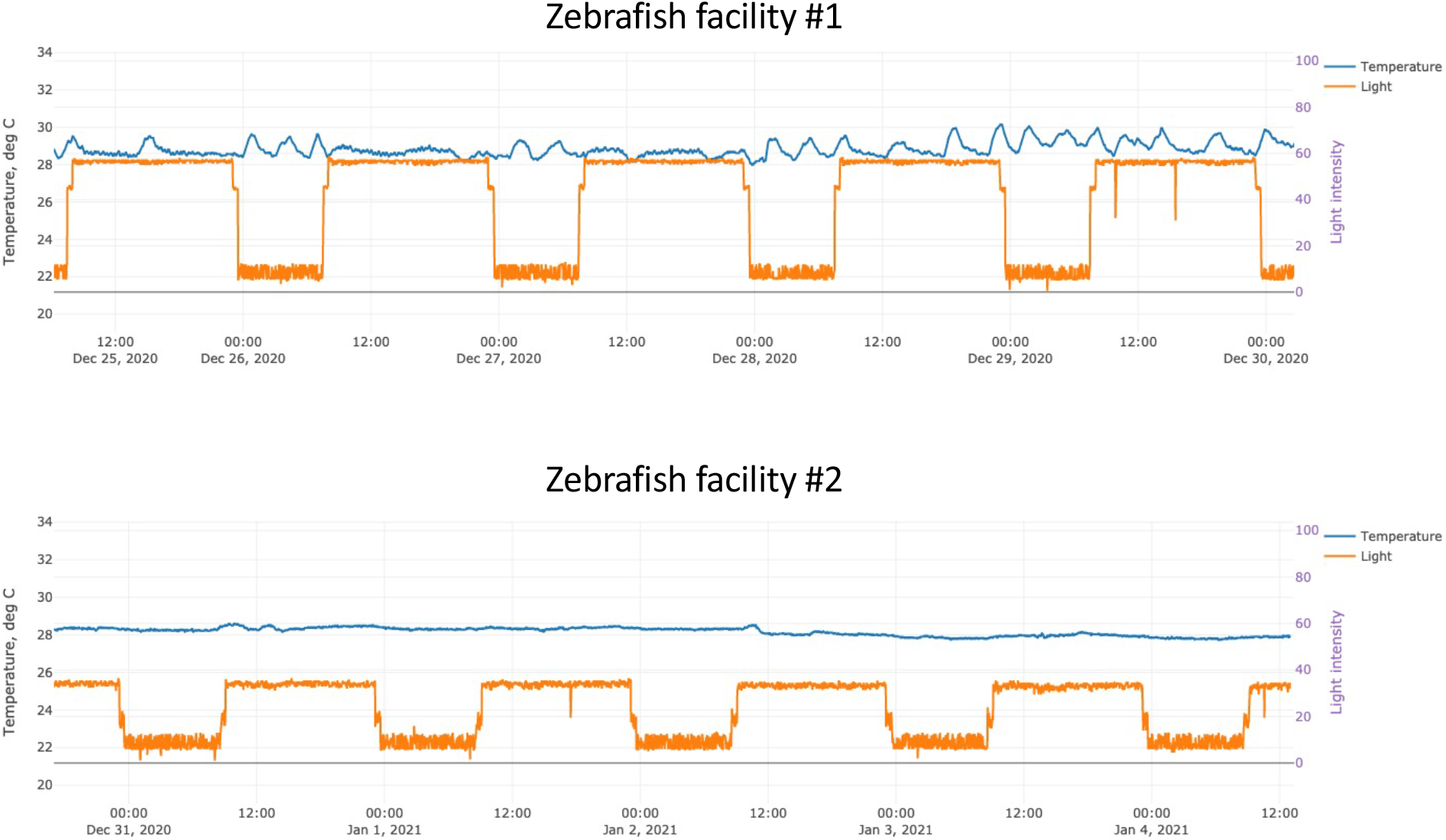
Continuous recordings of temperature and light levels from two zebrafish facilities. Data was collected using the same Raspberry Pi device with web interface. Light level has two values above dark: white lights that were turned on from 9 am until 11 pm, and red lights that were turned on for 30 minutes immediately before and after white lights were turned on during the day. The two facilities are located in different buildings on the same campus. Data was collected in 1 minute intervals.

**Supplemental Figure 2.**
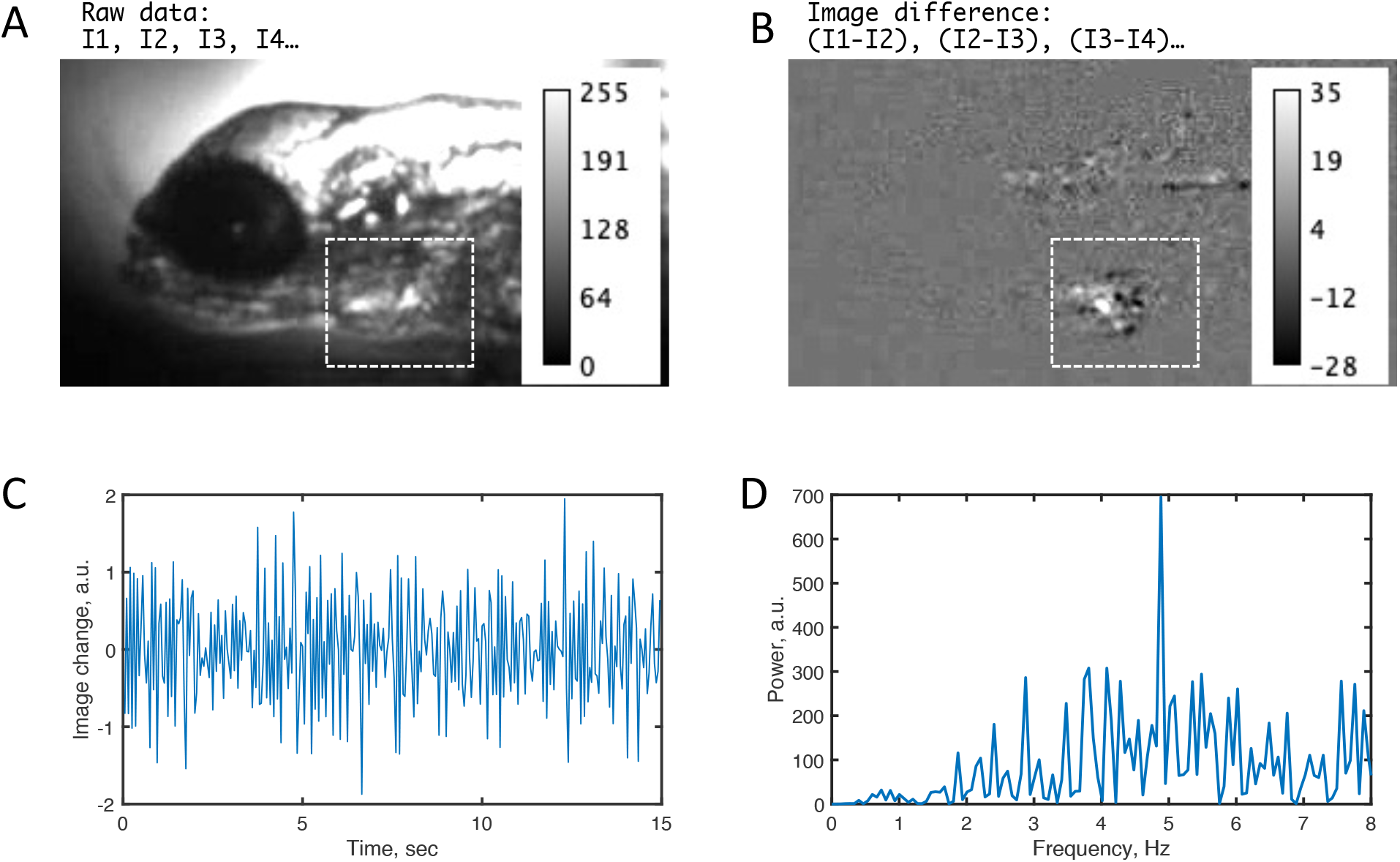
Heart rate extraction from video recording of larval zebrafish heart. (**A**) Trans-illuminated microscopy images of a zebrafish heart were acquired at 20 frames per second as a sequence of frames (I1, I2, I3…). The white box outlines the location of the heart. (**B**) Image difference between consecutive frames was generated using the Image Calculator function of Fiji. (**C**) Average image intensity in the region of interest (ROI) around the heart of the image-difference data was extracted. (**D**) Fourier-transformed data shows a peak at 5 Hz.

